# Rearrangement of chromosome spatial organization in mouse zygotes using optical tweezers does not affect preimplantation development

**DOI:** 10.64898/2026.10.26.761794

**Authors:** Corentin Mollier, Ozan Kastas, Laure de Chancel, Julia Roensch, Lorena Kolar-Znika, Anne de Cian, Maxime Dahan, Daniele Fachinetti, Antoine Coulon, Jean-Léon Maître

## Abstract

The spatial arrangement of chromosomes is thought to influence key processes such as gene expression or chromosome segregation. However, the contribution of chromosome spatial organization in physiological contexts remains disputed. Here, we use optical tweezers to displace nucleoli and physically rearrange the nuclear contents of mouse zygotes. Monitoring the preimplantation development of zygotes with physically shuffled nuclei reveals that cell dynamics, gene expression and overall blastocyst formation are not affected. This suggests that in mouse zygotes, the spatial organisation of chromosomes is not critical for blastocyst development.

## Introduction

The spatial organization of the genome is thought to regulate key biological processes. For example, the position of a gene may influence its expression, with examples of genes shutting down when tethered to the nuclear envelope (1, 2), while other studies contradict this picture (3). Similarly, chromosome mis-segregation during mitosis correlates with their position during interphase and forcing a chromosome to bind the nuclear envelope increases its likelihood to missegregate (4). In vivo, major genetic perturbations of chromosome organization by fusions of telomeres to telomeres or centromeres yields mild or limited impact on mouse development, depending on the chromosomes involved (5, 6). Most often, functional perturbation of 3D chromosome organization relies on genetic or chemical perturbations, which, although powerful and elegant, may introduce confounding effects with limited scope on their physiological impact. Therefore, testing the relevance of chromosome spatial organization would benefit from alternative methods compatible with in vivo settings.

During mouse preimplantation development, chromosome position was proposed to carry epigenetic memory and influence gene expression until the 4-cell stage (7). Also, in human zygotes, abnormal chromosome positioning within parental pronuclei before mitosis correlates with chromosome mis-segregation and blastocyst formation (8). However, these correlations remain untested.

## Results

To investigate the contribution of genome spatial organisation during mouse preimplantation development, we performed optical tweezers manipulation of zygote nucleolar precursor bodies (NPBs) in otherwise untreated embryos (Video 1). NPBs are RNA–protein phase-separated organelles coated with pericentromeric and telomeric regions (9) that exhibit strong refractive-index contrast with the surrounding nucleoplasm, allowing us to reliably manipulate them using optical tweezers (Fig. 1A). Depending on the number of NPBs present, we were able to perform three distinct types of perturbation (Fig. 1B, Video 2). 1) Rotation: When only one NPB was present, we used the optical trap to perform 3-8 complete revolutions. 2) Fusion: When multiple NPBs were present, we used multiple traps to bring them into contact, triggering their fusion. 3) Twisting: When multiple NPBs were present, we first fused a subset of them and then performed a twisting motion by alternately displacing the two remaining NPBs for 3-5 complete revolutions. NPBs fusion produces the most irreversible rearrangement, as chromosome regions that were previously associated with distinct NPBs are brought into closer proximity. Rotation produces displacements over larger spatial scales, as NPBs undergo several complete revolutions within pronuclei. Twisting combines both types of rearrangements for maximal perturbation of nuclear organization. Throughout the study, we used a control manipulation, referred to as Control OT clamp, in which the laser is applied to the centre of NPBs without displacing them.

**Fig. 1.**
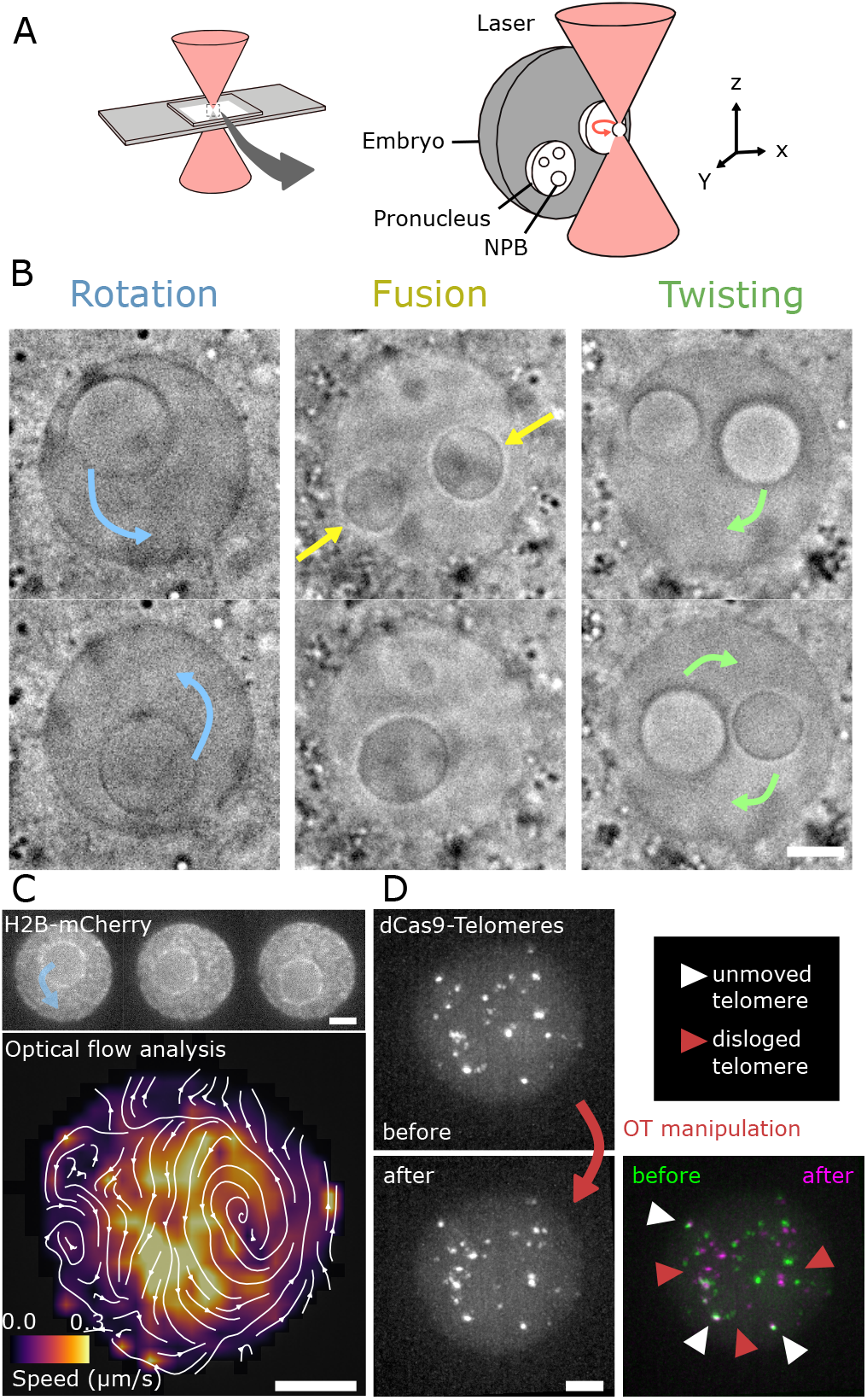
Optical tweezer-mediated manipulation of nuclear organisation. A) Schematic diagram of the experimental approach used to physically perturb nuclear organization. Left, embryo mounting. Right, cross-section of the embryo and displacement of nucleolus precursor bodies (NPBs) using optical tweezers. B) Brightfield images illustrating the types of manipulations of NPBs. For rotation and twisting, 2-8 and 3-5 revolutions are performed, respectively. Scale bar, 5 µm C) H2B-mCherry movement associated with a rotation (top) and corresponding optical flow analysis (bottom). Scale bar, 5 µm. D) dCas9-HaloTag with a sgRNA labeling telomeres before (top left, green) and after (bottom left, magenta) perturbation (twisting) and their overlay (bottom right). Scale bar, 5 µm.

To qualitatively assess the perturbation of nuclear organization, we used optical tweezers on H2B-mCherry expressing embryos, which exhibit enriched signal around NPBs due to their association with pericentromeric and telomeric regions (9) and a granular pattern throughout the nucleoplasm (Fig. 1C, Video 3). During NPB displacement, NPB-associated chromatin is displaced together with the NPB while nucleoplasmic foci are extensively rearranged, indicating that chromatin is displaced during the manipulation. Because individual H2B foci are difficult to track, we further visualised the displacement of individual genomic loci by microinjecting fluorescently labelled dCas9 with a sgRNA targeting telomeres (Fig. 1D, Video 4). Three-dimensional imaging before and after confirmed that optical tweezers manipulation induces spatial rearrangements of telomeres beyond those at NPB surface.

After perturbation, embryos were cultured individually until the blastocyst stage. Control embryos that remained in the incubator developed more frequently to the blastocyst stage than those mounted in the manipulation chamber, exposed to the imaging and tweezer light (Fig. 2A). However, we observed no difference in blastocyst formation between perturbed and control embryos (Fig. 2A), regardless of the types of perturbations (fusion, rotation or twisting). Therefore, for the remainder of our analysis, we group these perturbations together.

**Fig. 2.**
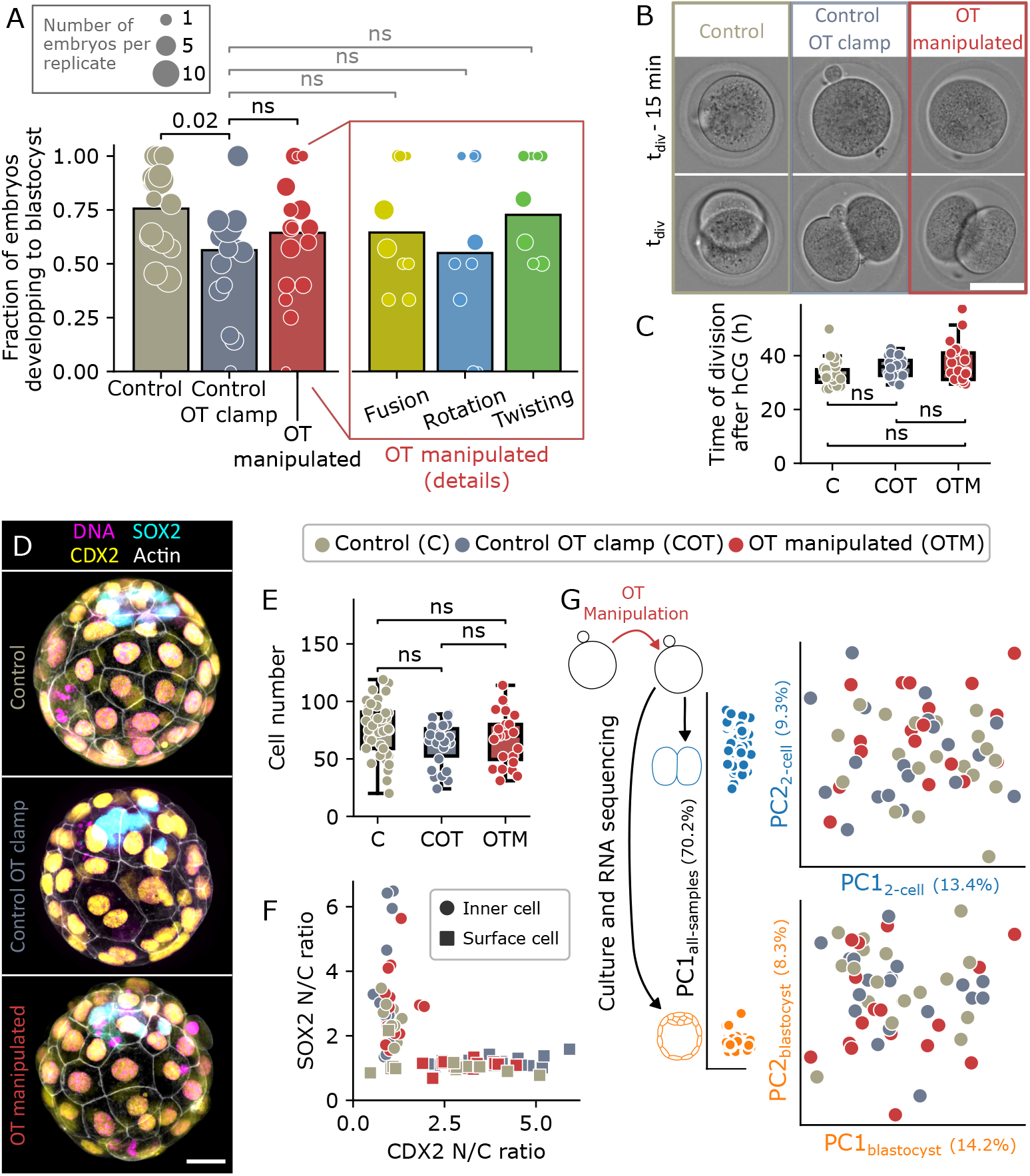
Spatial rearrangement of the genome at the zygote stage does not impact preimplantation development. A) Blastocyst turnout rate at E4.5. Each dot represents one replicate and their size corresponds to the number of embryos within each replicate. Boxes correspond to the total fraction that turned to blastocyst for each group: 139 Control, 99 Control OT clamp, 31 Fusion, 15 Rotation, 19 Twisting embryos from at least 8 replicates. Two-proportion Z-test, ns for not significant. B) Time-lapse imaging of zygotes to 2-cell stage division (t_div_). Scale bar, 50 µm. C) Time between human chorionic gonadotropin (hCG) injection and 2-cell division. 30 Control, 18 Control OT clamp and 24 OT manipulated embryos from 4 replicates. Statistical significance was determined by Mann-Whitney U test, ns not significant. D) Blastocysts immunostaining showing CDX2 (yellow), SOX2 (cyan), DAPI (magenta), phalloidin (white). Scale bar, 20 µm. E) Blastocyst cell number. 38 Control, 26 Control OT clamp and 23 OT manipulated embryos from 8 replicates. Statistical significance was determined by Mann-Whitney U test, ns not significant. F) Nuclear to cytoplasmic (N/C) ratio of CDX2 and SOX2 for individual cells. 5 inner and 5 surface cells for each embryo with 3 embryos per condition. G) Principal-component analysis (PCA) of RNA sequencing data between both stages (left), or between treatments at 2-cell (top right) and blastocyst (bottom right) stages. PCA was performed independently for each dataset. Percentage of the variance explained indicated in each axis. 2-cell : 20, 19 and 18 embryos; blastocyst: 18, 18 and 17 embryos for Control, Control OT clamp and OT manipulated, respectively, from 4 replicates each. P-values were adjusted for multiple comparisons using the Bonferroni method.

To inquire into potential effects on cell dynamics and/or gene expression during preimplantation development, we first asked whether the timing of the first cleavage was affected, as delays could indicate compensatory processes, such as the time required to fix genome spatial arrangement or activation of the spindle assembly checkpoint during mitosis (4, 8). To address this, we performed live imaging and quantified the timing of cell division relative to hCG injection (Fig. 2B-C). We detected no difference in cleavage timing, suggesting that physical perturbation of chromosome spatial arrangement has no effect on checkpoints activation or early division dynamics. Next, we analyzed the composition of blastocysts. Immunostaining revealed no change in the total cell number after perturbation of chromosome spatial arrangement at the zygote stage (Fig. 2D-E), confirming the overall lack of effect on cell cycling or cell division throughout preimplantation development.

To evaluate correct patterning of blastocysts after perturbation of chromosome spatial arrangement, we used immunostaining of CDX2 and SOX2, markers of the trophectoderm and inner cell mass, respectively (Fig. 2D). Quantifying the nuclear-to-cytoplasmic ratios of individual inner and outer cells (Fig. 2F), both markers showed their expected specificity in both control and perturbed embryos. Beyond the patterning of a couple of markers, we also assessed the potential impacts of perturbation of chromosome spatial arrangement at the zygote stage on overall transcription. We performed single-embryo RNA sequencing at the 2-cell and blastocyst stages following perturbations. Principal component analysis shows clear segregation of 2-cell and blastocysts expression profiles, but fails to separate the transcriptomes of control and perturbed embryos (Fig. 2G). Furthermore, differential gene expression analysis identified no significant differences between groups (significance threshold of 0.05 for Wilcoxon test with Benjamini–Hochberg multiple testing correction). This further confirms that disrupting chromosome spatial arrangement at the zygote stage does not affect cell dynamics or gene expression during blastocyst development.

## Discussion

In this work, we present a novel non-invasive method to induce spatial rearrangements of the genome in vivo through physical perturbations using optical tweezers. We apply this method to mouse zygotes, for which chromosome organization has been proposed to influence gene expression (7). Our results show that physically rearranging nuclear content at the zygote stage does not impair subsequent development, arguing against a critical role for chromosome spatial organization at this stage. Instead, this suggests that genome organization at this stage is plastic and/or rapidly re-established following perturbation.

This finding is particularly relevant in the context of zygotic genome activation (ZGA). In mice, ZGA occurs in two phases: a minor wave at the zygote stage, coinciding with DNA replication, followed by a major wave at the 2-cell stage. The minor ZGA has been shown to be essential for proper development (10). Given the timing of our perturbations, it is likely that we predominantly affect this minor wave. Nevertheless, the absence of detectable transcriptional or developmental defects suggests that minor ZGA is largely permissive to large-scale changes in 3D genome organization.

Together, our results support a model in which early embryonic genome function operates independently of fully established higher-order chromatin architecture, with large-scale 3D chromosome organization not bearing significant functional relevance at this stage of development.

## Material and Methods

### Embryo recovery and culture

All mouse work is performed under the supervision and permission of the institutional veterinarian and in agreement with governmental approval (APAFIS number 39490-2022111819233999 v2). The animal facilities are operated according to international animal welfare rules. Mice are used from 4 weeks old on. (C57BL/6xC3H) F1 hybrid strain is used as wild-type. To visualize chromatin R26-H2BmCherry CDB0239K mice are used (11).

Superovulation of female mice is induced by intraperitoneal injection of five international units of pregnant mare’s serum gonadotropin (Ceva, Syncropart), followed by intraperitoneal injection of five international units of human chorionic gonadotropin (MSD Animal Health, Chorulon) 44–48 h later. Embryos are recovered at E0.5 by tearing up the ampulla in 37°C homemade KSOM HEPES (12) (NaCl (Honeywell Research Chemicals, Fluka Cat31434), KCl (Sigma, BioReagent P540 LifeGlobal, ZEKS-050), KH_2_PO_4_ (Avantor Science Central, 26936.293), Na Lactate (Sigma, L4263), Na pyruvate (Sigma P4562), CaCl_2_ 2H_2_O (FisherScientific, C79-500), MgSO_4_ 7H_2_O (Sigma, 230391), Na_4_EDTA 4H_2_O (Sigma, E-5391) MEM essential AA solution (FisherScientific GIBCO, #11130-036), non-essential AA solution (FisherScientific GIBCO, #11140-035), L-Glutamine (FisherScientific GIBCO, #25030-032), D(+) glucose. (Sigma, G-6152), NaHCO_3_ (Sigma, S-5761), Penisilin and Streptomycin (FisherScientific GIBCO, 15140122), HEPES (Sigma, H4034)).

Embryos are placed for 10 s with 37°C 0.3 mg/mL hyaluronidase (Sigma, H4272-30MG) to remove surrounding cumulus cells and then washed in KSOM HEPES. Finally, embryos are transferred in 10 µL drops of KSOM supplemented with 1 mg/mL BSA (Sigma Aldrich M8410) covered in oil (Sigma, M8410 or Acros Organics) at 37°C and 5% CO2 in a humidified atmosphere.

Embryos are handled using an aspirator tube (Sigma, A5177-5EA) equipped with a glass pipette pulled from glass micropipettes (Blaubrand intraMark or Warner Instruments).

### Immunostainings

Embryos are fixed in 2% PFA (Euromedex, 2000-C), washed in PBS, and permeabilized in 0.01% (zygotes) or 0.1% (blastocysts) Triton X-100 (Euromedex, T8787) in PBS (PBT) at room temperature before being placed in blocking solution (PBT with 3% BSA) at 4°C for 2–4 hour. Primary antibodies were applied in blocking solution at 4°C overnight. After washes in PBT at room temperature, embryos were incubated with secondary antibodies, DAPI, and/or phalloidin in blocking solution at room temperature for 1 hour. Embryos were washed in PBT and imaged immediately after.

#### Primary antibodies

CDX2 (rabbit; dilution 1:200; Abcam, ab76541) and SOX2 (rat; dilution 1:100; Invitrogen, 14-9811-82).

#### Secondary antibodies and fluorescent dyes

4’,6-diamidino-2-phenylindole (DAPI; dilution 1:100; Invitrogen, D1306); Alexa Fluor 488 phalloidin (dilution 1:200; Fisher Scientific, 10125092); goat anti-Rabbit IgG (H+L) Cross-Adsorbed Secondary Antibody, Alexa Fluor 546 (dilution 1:200; Fisher Scientific, A11010); and goat anti-Rat IgG (H+L) Cross-Adsorbed Secondary Antibody, Alexa Fluor 633 (dilution 1:200; Fisher Scientific, A21094).

### dCas9-telomere microinjections

#### Protein purification

dCas9m4-NLS-HaloTag-NLS is produced in E.coli strain Rosetta 2(DE3) (Novagen, Sigma-Aldrich) and purified from 1 L of 2X YT (MP Biomedicals) medium supplemented with Carbenicillin 50 µg/mL(MP Biomedicals), Chloramphenicol 34 µg/mL (Sigma-Aldrich) and 1% Glucose after induction with 0.3 mM IPTG at OD600 = 0.7 and growth at 19°C for 16 h with orbital shaking at 200 rpm. Bacterial pellet is washed with 1X PBS (Gibco) and lysed by sonication (Vibracell 75186) for 10 min at 50% Amplitude (15 s ON/ 15 s OFF) in buffer HisA containing 25 mM Tris-OAc pH8, 500 mM NaCl, 20 mM Imidazole, 1 mM TCEP, supplemented with 1.5X Protease Inhibitor Ultra EDTA free (Roche). Insoluble material is centrifuged at 13000 g at 4°C for 30 min and supernatant was applied on a HisTRAP crude FF 5 mL (Cytiva), equilibrated in Buffer HisA. His column is washed with 100-200 mL of buffer HisA and eluted with step or gradient of Buffer His B containing 25 mM TrisOAc pH8, 500 mM NaCl, 250 mM Imidazole, 10% Glycerol and 1 mM TCEP. Eluted protein is concentrated on Amicon 100 kDa (Merck Millipore) and buffer exchanged by adding 4 volumes of Buffer ExA containing 20 mM HEPES-NaOH pH7.5, 100 mM KCl, 10% Glycerol and 1 mM TCEP. Protein is applied on HiTrap SP sepharose HP 1 mL (Cytiva) preequilibrated with Buffer ExA and gradient eluted between 33 to 50% of Buffer ExB (20 mM Hepes-NaOH pH7.5, 1 M KCl, 10% Glycerol, 1 mM TCEP). Protein is concentrated to 300 µL on Amicon 100 kDa (Merck Millipore) and applied to a gel filtration column Superdex 200 Increase 3.2/300 (Cytiva) pre-equilibrated with 5.5% of Buffer ExB in Buffer ExA. The fraction containing dCas9Halo protein is concentrated to 37-42 µM aliquoted and flash frozen at -80°C.

#### Protein sequence

MDKKYSIGLAIGTNSVGWAVITDEYKVPSKKFKVLG DRHSIKKNLIGALLFDSGETAEATRLKRTARRRYT RRKNRICYLQEIFSNEMAKVDDSFFHRLEESFLVEED KKHERHPIFGNIVDEVAYHEKYPTIYHLRKKLVDSTD KADLRLIYLALAHMIKFRGHFLIEGDLNPDNSDVDKL FIQLVQTYNQLFEENPINASGVDAKAILSARLSKSRRL ENLIAQLPGEKKNGLFGNLIALSLGLTPNFKSNFDLAE DAKLQLSKDTYDDDLDNLLAQIGDQYADLFLAAKNL SDAILLSDILRVNTEITKAPLSASMIKRYDEHHQDLTL LKALVRQQLPEKYKEIFFDQSKNGYAGYIDGGASQEE FYKFIKPILEKMDGTEELLVKLNREDLLRKQRTFDNG SIPHQIHLGELHAILRRQEDFYPFLKDNREKIEKILTFRIPYYVGPLARGNSRFAWMTRKSEETITPWNFEEVVDK GASAQSFIERMTNFDKNLPNEKVLPKHSLLYEYFTVY NELTKVKYVTEGMRKPAFLSGEQKKAIVDLLFKTNR KVTVKQLKEDYFKKIECFDSVEISGVEDRFNASLGTY HDLLKIIKDKDFLDNEENEDILEDIVLTLTLFEDREMIE ERLKTYAHLFDDKVMKQLKRRRYTGWGRLSRKLIN GIRDKQSGKTILDFLKSDGFANRNFMQLIHDDSLTFK EDIQKAQVSGQGDSLHEHIANLAGSPAIKKGILQTVK VVDELVKVMGRHKPENIVIEMARENQTTQKGQKNSR ERMKRIEEGIKELGSQILKEHPVENTQLQNEKLYLYY LQNGRDMYVDQELDINRLSDYDVAAIVPQSFLKDDSI DNKVLTRSDKARGKSDNVPSEEVVKKMKNYWRQLL NAKLITQRKFDNLTKAERGGLSELDKAGFIKRQLVET RQITKHVAQILDSRMNTKYDENDKLIREVKVITLKSK LVSDFRKDFQFYKVREINNYHHAHDAYLNAVVGTA LIKKYPKLESEFVYGDYKVYDVRKMIAKSEQEIGKAT AKYFFYSNIMNFFKTEITLANGEIRKRPLIETNGETGEI VWDKGRDFATVRKVLSMPQVNIVKKTEVQTGGFSK ESILPKRNSDKLIARKKDWDPKKYGGFDSPTVAYSVL VVAKVEKGKSKKLKSVKELLGITIMERSSFEKNPIDFL EAKGYKEVKKDLIIKLPKYSLFELENGRKRMLASAGE LQKGNELALPSKYVNFLYLASHYEKLKGSPEDNEQK QLFVEQHKHYLDEIIEQISEFSKRVILADANLDKVLSA YNKHRDKPIREQAENIIHLFTLTNLGAPAAFKYFDTTI DRKRYTSTKEVLDATLIHQSITGLYETRIDLSQLGGDG GGSGTSSPKKKRKVGIGSGSNGSSGAEIGTGFPFDPHY VEVLGERMHYVDVGPRDGTPVLFLHGNPTSSYVWR NIIPHVAPTHRCIAPDLIGMGKSDKPDLGYFFDDHVRF MDAFIEALGLEEVVLVIHDWGSALGFHWAKRNPERV KGIAFMEFIRPIPTWDEWPEFARETFQAFRTTDVGRK LIIDQNVFIEGTLPMGVVRPLTEVEMDHYREPFLNPV DREPLWRFPNELPIAGEPANIVALVEEYMDWLHQSPV PKLLFWGTPGVLIPPAEAARLAKSLPNCKAVDIGPGL NLLQEDNPDLIGSEIARWLSTLEISGEPTTGGRLPKKK RKVPRGGGGSHHHHHH*

#### HaloTag and sgRNA complexation

dCas9 protein is labeled in vitro with JF549 HaloTag ligand (Promega, GA1110) in a 1:3 molar ratio. Labeling reaction is carried out in a buffer containing 150 mM KCl, 20 mM HEPES, 10% glycerol and 1 mM TCEP (Sigma, ref 646547) at pH 7.5 and 24°C for 2 h. Excess ligand is removed using Micro Bio-Spin P-30 Gel Columns (Bio-Rad, 7326250). The buffer of the column is exchanged with a labeling reaction buffer. Protein concentration is measured using a nanodrop. Labeled dCas9 protein is snap frozen in liquid nitrogen and stored in a labeling reaction buffer at -80°C. Thawed JF549 labeled dCas9 protein and sgRNA (Merck, VC40007. Sequence : GUUAGGGUUAGGGUUAGGGUUA. (13)) are assembled in a 1:30 molar ratio with 0.5 µM of the dCas9 solution and 15 µM of sgRNA targeting telomeres in injection buffer (5 mM Tris–HCl pH = 7.4, 0.1 mM EDTA) for 10 min at 37°C.

#### Microinjection

For dCas9 microinjections, glass capillaries (Harvard Apparatus glass capillaries with 780 µm inner diameter) are pulled using a P-97 Flaming Brown needle puller (Sutter Instrument) and microforge (Narishige, MF-900) to build a holding pipette and an injection needle. The resulting injection needles are filled with 3 µL of the dCas9 sgRNA solution. The filled needle is positioned on a micromanipulator (Narishige MMO-4) and connected to a positive pressure pump (Eppendorf FemtoJet). Embryos are placed in KSOM HEPES supplemented with BSA drops covered with mineral oil under microscope (Leica DMi8). Zygotes are injected while holding with a holding pipette connected to a Micropump Cell-Tram Oil.

### Microscopy

#### Spinning disc microscope with optical tweezers

For manipulations with optical tweezers, embryos are mounted in KSOM HEPES using a custom 500 µm steel spacers (based on a design kindly provided by the lab of Pere Roca-Cusachs) in between glass coverslips (VWR, 631-0125). A multimodal platform is used that combines spinning-disc confocal microscopy and optical micromanipulation. Briefly, an optical micromanipulation unit (SENSOCELL, IMPETUX) modulated with a pair of acousto-optic deflectors is coupled to the rear epifluorescence port of an inverted research microscope (Nikon Ti2 Eclipse). The optical traps are created at the focal plane of a ×60/1.2NA water-immersion objective (Nikon). Dynamic control of the optical traps is performed with the manufacturer software (LightACE 1.6.2, IMPETUX). Simultaneous optical trapping is performed by time sharing the laser spot at 25 kHz. An around 130 mW trap is used (1 W of output laser power at 1064 nm). Power and position calibration are run systematically before experiments. The microscope is equipped with an incubation chamber to keep the sample at 37°C (Emblem).

Imaging is performed using an inverted Nikon Eclipse Ti2 microscope equipped with an Oxxius laser source providing excitation wavelengths of 488, 561, and 638 nm, each with a nominal output power of 100 mW. The system is equipped with a Yokogawa CSU-X1 spinning-disk confocal unit with a 50 µm pinhole disk and a Hamamatsu ORCA-Flash4.0 LT Plus camera. Image acquisition was controlled using Micro-Manager 2.0.

#### Spinning disc microscope

For 3D confocal imaging of immunostainings, an inverted Zeiss Observer Z1 microscope with a CSU-X1 spinning disk unit (Yokogawa) is used. Excitation is achieved using 405, 488, 561, and 642 nm laser lines through a ×63/1.2 C Apo Korr water immersion objective. Emissions are collected through 450/ 50 nm, 525/50 nm, 595/50 band pass or 610 nm low pass filters onto an ORCA-Flash 4.0 camera (C11440, Hamamatsu). This microscope is operated using the Metamorph 7.10.4.407 software.

#### Widefield microscope

For time-lapse imaging of embryo development to the 2-cell stage, embryos are placed in 5 cm glass-bottom dishes (MatTek) under a Celldiscoverer 7 (Zeiss) equipped with a x20/0.95 objective and an ORCA-Flash 4.0 camera (C11440, Hamamatsu). Embryos are imaged every 15 min at three focal planes positioned 10 µm apart. Embryos are kept in a humidified atmosphere supplied with 5% CO2 at 37°C.

### Optical tweezers manipulations

Embryos are mounted in KSOM HEPES using a custom 500 µm steel spacers (based on a design kindly provided by the lab of Pere Roca-Cusachs) in between glass coverslips (VWR, 631-0125). Simultaneous optical trapping is performed by time sharing the laser spot at 25 kHz using 130 mW per trap (1 W of output laser power). To disrupt nuclear organisation, we either perform a rotation, fusion or twisting manipulation. The same manipulation is done on both pronuclei. Rotation consists in trapping a single NPB and dragging it throughout the equatorial plane of the pronucleus 3-8 times clockwise or counter clockwise. Rotation is systematically chosen when one of the pronuclei contains a single NPB. Fusion consists in the fusion of all NPBs into a single one by forcing NPBs together using two traps. Twisting consists in sequential rotational movements of two NPBs within the equatorial plane of the NPBs (3-5 complete revolutions). Twisting is often preceded by the fusion of several NPBs so that sufficient space is available within the pronuclei to perform the twisting motion.

To analyse the effect of NPB movements on chromatin distribution, NPB trapping is done on H2B-mCherry expressing embryos using the spinning disc configuration of the microscope.

To analyse the disruption of nuclear organisation, manipulations can be preceded and followed by live imaging of telomeres. Telomeres are imaged using the spinning disc configuration before optical tweezer perturbations are performed using the brightfield configuration.

Additional traps are sometimes placed in the cytoplasm to immobilize the embryo.

### Image analysis

#### Optical flow analysis

Optical flow is based on the Lucas-Kanade method that estimates the displacement of a sparse set of points between two consecutive frames, and is implemented in python. From H2B-mCherry confocal images, the nucleus is first segmented independently in each frame (Gaussian smoothing, 50^th^-percentile threshold, morphological closing, hole filling, largest connected component), and the resulting mask restricts where points are sought. Background signal (Gaussian blur, r = 10 px) is subtracted from each frame and minmax normalised. Foci inside the nucleus are identified with the Shi-Tomasi detector (i.e. the most textured locations of the chromatin with `cv2.goodFeaturesToTrack`, up to 150 points in the image, at least 7 px apart), and tracked from frame to frame with the Lucas-Kanade method. It then solves, by least squares, for the displacement that minimizes the intensity difference between the two frames, iteratively and over an image pyramid to capture larger motions (`cv2.calcOpticalFlowPyrLK`, 21 × 21 px window, 3 pyramid levels) and re-detected every 10 frames to compensate for lost points. Vectors are averaged on a spatial grid, interpolated, and displayed as streamlines.

#### Immunostaining analysis

SOX2 and CDX2 expression levels are measured using (14). For each embryo, 10 cells (five inner cells, five surface cells) are measured. For each cell, the signal intensity is measured in a representative 21 µm^2^ circular area of the nuclear equatorial plane and in a directly adjacent cytoplasmic area. Mitotic and apoptotic cells are excluded from analysis. We then calculate the corresponding nuclear to cytoplasmic ratios.

### Statistical analysis

In box plots, the central line indicates the median, boxes represent the interquartile range (25^th^–75^th^ percentiles), and whiskers extend to 1.5 times the interquartile range. Individual data points are overlaid. For blastocyst turnout rates, manual implementation in python of two-proportion z-test is used. For all other statistical analysis, Mann-Whitney utests are used from the statannotation package (15). P-values are systematically corrected for multiple testing using Bonferroni correction.

### RNA sequencing

#### Lysis

In each well of PCR strips tubes (ThermoFisher Scientific, AB-0266), 1 µL of lysis buffer containing 0.02 µL Triton (ThermoFisher Scientific, X100), 0.09 µL dCTP (100 mM, ThermoFisher Scientific) 0.03 µL Recombinant RNAse Inhibitor (40 U/µL, Takara Bio), 0.24 µL dNTP (25 mM each, ThermoFisher Scientific), 0.2 µL Betaine (5 M, ThermoFisher Scientific), 0.012 µL Dithiothreitol (100 mM, ThermoFisher Scientific), 0.092 µL oligo-dT30VN ((5 Bio-AAG CAGTGGTATCAACGCAGAGTACT30VN-3 ; Bio = C6-linker biotin; 100 µM, IDT) according to published Flash-seq protocol (Lightning Fast and Highly Sensitive Full-Length Single-cell sequencing using FLASH-Seq, (16)) and ultrapure RNase-free water (Fisher 10977049, Invitrogen UltraPure DNase/RNase-Free Distilled Water). Embryos are loaded individually using mouth pipetting on ice before being frozen at -80°C until further use.

#### Library preparation

PCR strips tubes are transferred to a Biorad Thermocycler C1000 Touch set to 72°C for 3 min. Then, 4 µL of RT-PCR mix is dispensed (0.8 µL Betaine (5 M, ThermoFisher Scientific), 0.05 µL of MgCl2 (1 M, Invitrogen), 0.24 µl DTT (100mM ThermoFisher Scientific), 0.1 µL of RRI (40 U/µL, Takara Bio), 2.5 µL of KAPA HiFi Hot-Start ReadyMix (2X, Roche), 0.05 µL Maxima H Minus reverse transcriptase (200 U/µL), 0.092 µL TSO (5’-/5Biosg/AAGCAGTGG TATCAACGCAGAGTNNNNNNNNrGrGrG-3’ ; 100 µM, IDT) and water). The strip tubes are gently vortexed and spun down. Then, using the Biorad Thermocycler C1000 the following program is applied: 50°C for 60 min; 98°C for 3 min; 24 cycles at 98°C for 20 s, 67°C for 20 s, 72°C for 6 min. cDNA quality and concentrations are assessed using the Qubit dsDNA HS Assay (Invitrogen) and TapeStation D5000 (Agilent). Samples are transferred into a twin.tec PCR Plate 384 LoBind (Eppendorf twin.tec PCR Plate 384 LoBind 0030129547). Tagmentation is performed on 150 pg/µL of cDNA per well (Nextera XT DNA Library Preparation Kit, Illumina), mixing 1 µL of Tagment DNA buffer 4X, 0.25 µL of Amplicon Tagment mix and nuclease-free water for a final volume of 2 µL. The plate is incubated at 55°C for 9 min 20 s. The reaction is inactivated with 0.5 µL of 0.1% SDS (Sigma-Aldrich) for 5 min at room temperature. Finally, the tagmented products are amplified by PCR (KAPA HiFi PCR kit, Roche), mixing 1 µL of 5X GC Buffer, 0.15 µL of dNTP mix (10mM each), 0.09 µL HiFi enzyme, 0.5 µL of pre-mixed Illumina UD Indexes (5 µM each, IDT) and nuclease-free water for a final volume of 5 µL. Then, using the Biorad Thermocycler C1000 the following program is applied: 72°C for 3 min; 95°C for 30 s; 15 cycles at 95°C for 10 s, 55°C for 30 s, 72°C for 30 s. All the libraries are pooled and purified successively twice (SPRIselect beads, Beckman Coulter) with a ratio of 0.8X. The final library quality and concentrations are assessed using the Qubit dsDNA HS Assay (Invitrogen) and TapeStation D5000 (Agilent).

#### Sequencing and quality control

For sequencing, the library is diluted to 1 ng/µL and 5nM maximum. Paired-end sequencing was performed using a NextSeq 2000 sequencer with an average of 3M reads per cell. Flashseq data were processed using the pipeline scRNA-SmartSeq3 v3.2.0 developed within the Single Cell Initiative of Institut Curie and available at https://doi.org/10.5281/zenodo.23099833. Briefly, reads were aligned to the Mouse (mm10) reference genome with STAR (v2.7.8a) and assigned to genes using FeatureCounts (v2.0.1).

#### RNA sequencing analysis

Single-cell RNA-seq data are processed using the Scanpy framework (version 1.12). Embryos are first filtered based on quality control metrics, retaining only embryos with at least 8000 detected genes and fewer than 10% of reads mapping to mitochondrial genes. Following filtering, total counts per embryos are normalized using library size normalization (`sc.pp.normalize_total`) and log-transformed (`sc.pp.log1p`). Highly variable genes are then identified using Scanpy’s default method, selecting the top 2000 genes (`sc.pp.highly_variable_genes`). Principal component analysis (PCA) is subsequently performed on the dataset using the selected highly variable genes (`sc.tl.pca`) to reduce dimensionality for downstream analyses. Principal component analysis (PCA) is consistent with previously published results (data not shown; (17)).

Differential gene expression analysis between embryo groups is carried out using the non-parametric Wilcoxon rank-sum test as implemented in Scanpy (`sc.tl.rank_genes_groups`, method = “wilcoxon”). Statistical significance is assessed using default parameters, and resulting p-values are adjusted for multiple testing using the Benjamini–Hochberg procedure.

## ACKNOWLEDGEMENTS

We thank the imaging platform of the Genetics and Developmental Biology Unit (PICT-IBiSA@BDD, member of the French National Research Infrastructure France-BioImaging ANR-10-INBS-04), the CurieCoreTech Custom Single Cell Omics and Recombinant Protein platforms and the animal facility of the Institut Curie for their invaluable help. We thank Carine Giovannangeli and Jean-Paul Concordet for their help setting up the dCas9-HaloTag. We thank the team of Pere Roca Cusachs for sharing the design of their mounting chamber and the workshop of UMR168 at Institut curie for its fabrication. We thank Impetux for their help with optical tweezer control. We thank Ricardo Henriques and the Henriques Lab for making the bioRxiv LaTeX template publicly available, which was used to format this manuscript. Research in the lab of J.-L.M. is supported by the Institut Curie, the Centre National de la Recherche Scientifique (CNRS) and is funded by grants from the the European Research Council Consolidator Grant ERC-2022-CoG 101087656, the Coup d’élan from the Fondation Bettencourt Schueller and the Agence Nationale de la Recherche ANR-23-CE13-0020. Development of dCas9-HaloTag labeling was funded by the ANR (ANR-18-CE12-0023-01; ANR-22-CE95-0003-01) and National Science Foundation (NSF 2210558).

## AUTHOR CONTRIBUTIONS

C.M. and J-L.M designed the research; C.M., L.d-C. and O.K. performed experiments; J.R. A.d-C., L.K-Z, M.D and A.C. designed, optimised and provided dCas9-Halo labelling reagents; C.M. analyzed the data; C.M., D.F., A.C. and J-L.M. interpreted the data. C.M. and J-L.M. wrote the manuscript with inputs from all authors.

## Supplementary Movie Legends

**Video 1:** Time lapse imaging of nuclear organisation perturbation using optical tweezers to fuse all nucleolar precursor bodies from both pronuclei of a mouse zygote. Crosses mark the position of optical traps. Scale bar, 10 µm.

**Video 2:** Time lapse imaging of the different perturbations of nuclear organisation using optical tweezers. Each panel corresponds to a distinct treatment in different embryos.

**Video 3:** Time lapse imaging of H2B-mCherry (left) upon perturbation with optical tweezers (rotation). Middle shows 150 detected features with light red corresponding particle detection and dark red to their tracking for 10 consecutive frames before a new detection step is initiated. Right shows quantification of the magnitude and direction of the corresponding flow. Fig. 1C corresponds to the cumulative analysis of frames 29-50.

**Video 4:** Time lapse imaging of embryos injected with dCas9-NLS-HaloTag-NLS + JF549 and a sgRNA labeling telomeres before, during and after perturbation with optical tweezers. First part: max-projections of the entire nucleus before perturbation, showing endogenous movements of telomeres. Second part: brightfield imaging of perturbation with optical tweezers (twisting). Third part: overlay of the last image before perturbation (green) and first image after perturbation (magenta). Fourth part: max-projections of the entire nucleus after perturbation, showing endogenous movements of telomeres.

